# Role and impact of the gut microbiota in a *Drosophila* model for parkinsonism

**DOI:** 10.1101/718825

**Authors:** Virzhiniya Feltzin, Kenneth H. Wan, Susan E. Celniker, Nancy M. Bonini

## Abstract

*Drosophila* is poised to be a powerful model organism for studies of the gut-brain axis due to the relative simplicity of its microbiota, similarity to mammals, and efficient methods to rear germ-free flies. We examined the gut-brain axis in *Drosophila* models of autosomal recessive parkinsonism and discovered a relationship between the gut microbiota and *parkin* loss of function. The number of live bacteria was increased approximately five-fold in the gut of aged *parkin* null animals. Conditional RNAi showed that *parkin* is required in gut enterocytes and not in neurons or muscle to maintain microbial load homeostasis. To examine the significance of gut microbiota, we reared germ-free *parkin* flies and discovered that removal of microbes in the gut improves the animals’ resistance to paraquat. Sequencing of 16S rDNA revealed microbial species with altered relative abundance in *parkin* null flies compared to controls. These data reveal a role for *parkin* activity in maintaining microbial composition and abundance in the gut, suggesting a relationship between *parkin* function and the gut microbiota, and deepening our understanding of *parkin* and the impacts upon loss of *parkin* function.

## Introduction

Current studies have uncovered a fascinating link between the gut microbiota and the brain (Mayer et al., 2014; Sharon et al., 2016). For instance, alterations in the gut microbiota have been shown to affect host neurotransmitter levels, and anxiety- and depression-like symptoms (Bravo et al., 2011; Wong et al., 2016). In addition, studies suggest that changes in the gut microbiota are correlated with the development and severity of diseases such as autism and Parkinson’s disease (Hsiao et al., 2013; Sampson et al., 2016; Scheperjans et al., 2015). As promising as these initial studies are, in-depth research into the link between microbes in the gut and disease of the brain is challenging given the complexity of the mammalian microbiota and the intricacies presented by mammalian models.

The genetics powerhouse of *Drosophila* has the potential to facilitate breakthrough studies of the gut microbiota and their relation to disease. The microbiome of the fly gut is simpler than that of mammals, with up to 20 species comprising more than 90 percent of all bacteria in the gut (Fink et al., 2013; Wong et al., 2013), allowing for powerful reductionist studies. Of the well-known residents of the fly gut, the genera *Lactobacillus* and *Enterococcus* are also commonly present in the human gut microbiome (Arumugam et al., 2011; Eckburg et al., 2005; Qin et al., 2010). One can rear germ-free flies efficiently and at lower cost compared to mammals, enabling experimental screens and studies that examine the impact of the gut microbiota on various disease models. The *Drosophila* microbiota are passed from parent to larvae through contamination of the embryonic shell (chorion), which the larvae consume after hatching. The larval microbiota develop as the growing larvae eat, until reaching a plateau at the third instar stage, and it is then eliminated during the pupal stage. Newly eclosed adult flies have a very low number of live bacteria in the gut, and the gut microbiota grow in number and evolve in composition as the animals age (Broderick and Lemaitre, 2012).

We sought to harness the potential of the fly with a screen to investigate the gut/brain axis in fly models of human disease. *Drosophila* disease models have contributed to crucial discoveries of disease mechanisms and etiology due to the wide array of available molecular genetic tools and the many conserved genes and pathways (Bier, 2005; Marsh and Thompson, 2006). We initiated our studies by measuring the gut microbial abundance in loss-of-function mutants for genes associated with recessive parkinsonism: *parkin (park), PTEN-induced putative kinase 1 (pink1)*, and *DJ-1*. It is thought that the main contribution of Pink1 and Parkin to development of PD is through a pathway in which both proteins work towards maintaining mitochondrial fidelity (Greene et al., 2003; Park et al., 2006). In healthy mitochondria, Pink1 is rapidly degraded, but mitochondrial damage and depolarization causes Pink1 to accumulate on the outer mitochondrial membrane (OMM) (Jin et al., 2010; Meissner et al., 2011; Narendra et al., 2010). Pink1 phosphorylates Parkin resulting in recruitment of Parkin to the mitochondria and activation (Kane et al., 2014; Kazlauskaite et al., 2014; Kondapalli et al., 2012; Koyano et al., 2014; Shiba-Fukushima et al., 2012; Shiba-Fukushima et al., 2014), eventually leading to engulfment of the damaged mitochondrion (Sarraf et al., 2013). Parkin has also been shown to regulate mitochondrial fission and fusion, protect against intracellular bacterial pathogens, and together with Pink1 play a role in intestinal stem cell proliferation (Deng et al., 2008; Manzanillo et al., 2013; Park et al., 2006; Poole et al., 2008). DJ-1 senses oxidative stress through oxidation of its cysteine residues and protects the cell from the harmful effects of reactive oxygen species (Canet-Avilés et al., 2004; Hayashi et al., 2009; Martinat et al., 2004; Taira et al., 2004).

In examining the gut microbiota in these genes associated with parkinsonism, here we report a link between the gut microbiome and *parkin* mutant flies. We find the abundance of gut microbiota is increased in aged mutant *parkin* animals, and that the absence of gut microbiota ameliorates paraquat sensitivity in *parkin* animals. These findings suggest a bidirectional relationship between the gut microbiota and *parkin* gene function that affects the severity and progression of the gene mutation effects.

## RESULTS

### Microbial abundance is increased with age in *parkin* null animals

To explore the idea of interactions between *Drosophila* models of neurodegenerative disease and disturbances in the gut microbiota, we measured microbial abundance in the autosomal recessive parkinsonism models *parkin*^*1*^, *pink1*^*B9*^, and a double knockout for the two *DJ1* homologs in *Drosophila, DJ-1α* and *DJ-1β* (DJ-1 DKO). An abnormally high or low number of live bacteria in the gut indicates disruption of microbial homeostasis. Microbial abundance was quantified by dissecting the gut, homogenizing it through bead-beating, and spreading the homogenate in serial 10-fold dilutions on MRS-agar plates, a medium commonly used to rear the gut-associated microbes of *Drosophila* (Guo et al., 2014). The number of colonies that grew on the plates was counted and used to calculate the Colony Forming Units (CFU), representative of the number of live bacteria in the gut. We used males of ages 3d (young flies with a sparse microbiome) and 20d (older flies with a well-established abundant microbiome).

Consistent with previous findings (Guo et al., 2014),(Broderick et al., 2014), young flies had few living bacteria in the gut (∼10^3^), and this number rose steeply in older flies (∼10^5^) (Fig. 1a). There was no difference in microbial load between control flies and any of the parkinsonism gene models at 3d. At 20d, however, we observed a significant increase in the number of live microbes per gut of *parkin* null flies compared to control animals (∼10^6^) (Fig. 1a). Surprisingly, *pink1* and *DJ-1* mutant animals did not show a significant microbial load increase, even though Parkin and Pink1 are thought to regulate mitochondrial homeostasis and shape dynamics through the same pathway (Pickrell and Youle, 2015). This indicated a disturbance in the gut microbiota of *parkin* mutants, and that Parkin may play this role independently of Pink1.

**Figure 1.**
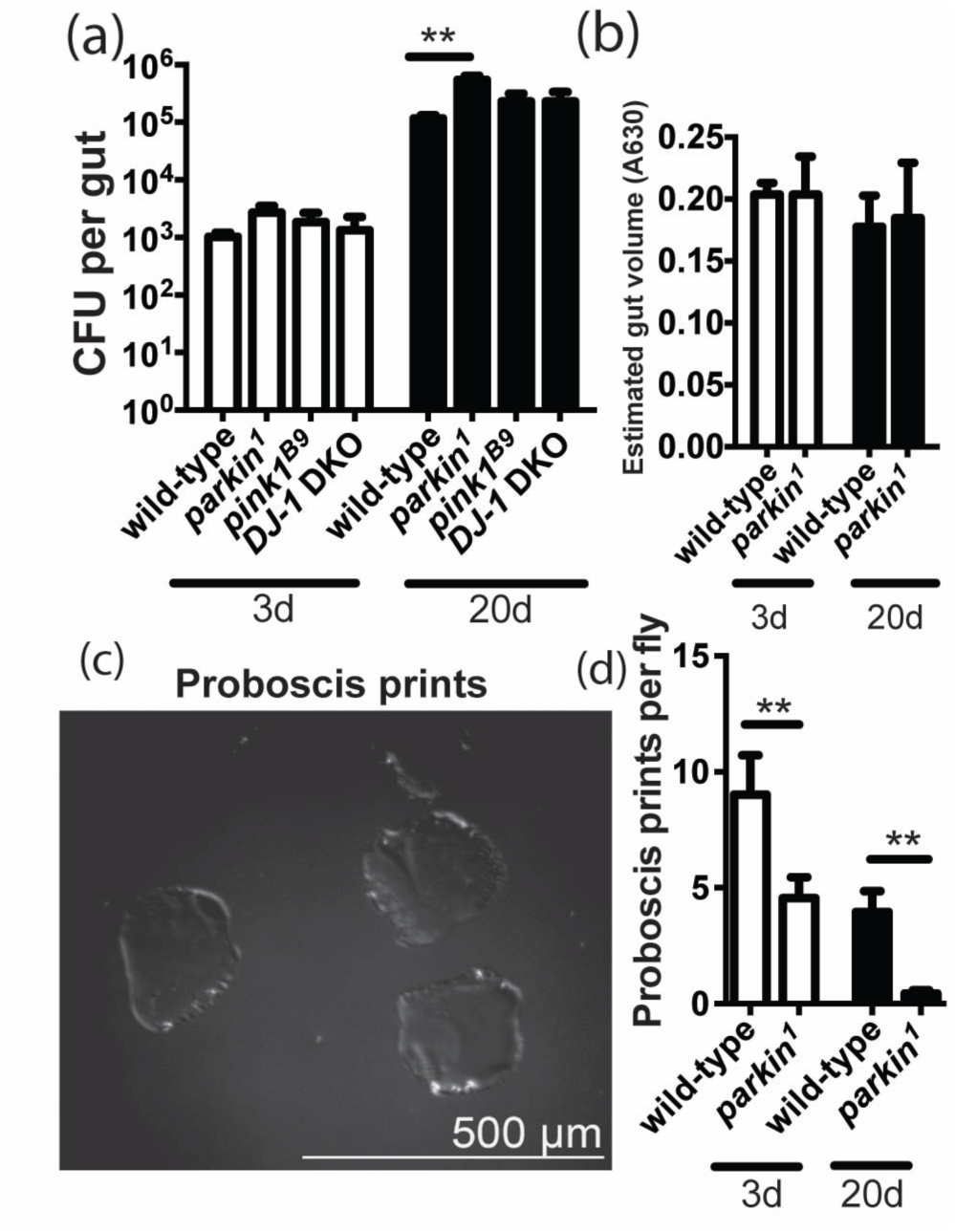
*parkin* mutants exhibit an elevated microbial load with age. (**A**) Microbial load of wild-type (*w*^*1118*^) males and male mutants for parkinsonism-associated genes at ages 3d and 20d. Dissected and homogenized individual guts were serially diluted and a fraction of the diluted homogenate was spread on MRS-agar plates. Colonies grown were counted and used to calculate the colony forming units (CFU) per gut. The experiment was repeated in four independent biological replicates of six individual guts each per age and genotype. DJ1 DKO stands for DJ1 double knockout: *DJ-1α*^*δ72*^; *DJ-1β*^*δ93*^. **p<0.01, ANOVA for significance, followed by Tukey’s post-test. Comparisons not marked with a double asterisk (**) are not statistically significant. **(B)** Blue-dye feeding assay to measure volume of food in the gut of wild-type (*w*^*1118*^) and *parkin*^*1*^ mutant males at 3d and 20d. Flies were placed on food containing 2.5% w/v FD&C blue dye #1 for 48 hr. Five guts per genotype/age group were dissected in PBS, homogenized, and the absorbance at 630 nm was measured. The experiment was repeated in three independent biological replicates. n.s. not significant, ANOVA followed by Tukey’s post-test. **(C)** Example image of prints left by the fly proboscis on a 1% gelatin-, 5% sucrose-coated slide. **(D)** Proboscis print assay to measure the rate of feeding of wild-type (*w*^*1118*^) and *parkin*^*1*^ mutant males at 3d and 20d. Animals were enclosed in individual chambers on top of a 1% gelatin-, 5% sucrose-coated slide and incubated for 20 min without disturbance. The number of proboscis prints left on the surface of the slide was counted. The experiment was repeated in ten independent biological replicates of ten individual flies each per age and genotype. **p<0.01, ANOVA followed by Tukey’s post-test.

We performed a series of control experiments to assess whether the increase in microbial load in *parkin* nulls was simply related to a change in eating or elimination from the gut. The rate of feeding was measured using proboscis print assays. Young and old wild-type and *parkin* male flies were placed individually on a microscope slide covered with sucrose-gelatin for 20 min(Edgecomb et al., 1994). As the fly ingests gelatin, the proboscis leaves a print on the surface of the slide, which was observed and scored using Differential Interference Contrast (DIC) microscopy (Fig. 1c). The number of proboscis prints left on the slide at the end of the assay reflects the rate of feeding. We determined that *parkin* flies eat significantly less than wild-type controls at 3d and 20d (Fig. 1d), suggesting the increase in microbial load cannot be due to increased feeding. To measure the volume of food in the gut, the flies were fed standard food supplemented with FD&C Blue Dye #1, then guts were dissected, homogenized, and the absorbance of the sample at 630nm was measured. The assay revealed no significant difference in gut volume between old and young *parkin* mutants and wild-type controls (Fig. 1b). Therefore, neither a higher rate of feeding, nor a larger volume of food in the gut explains the increased microbial load in the gut of *parkin* mutants.

Since the mutant animals eat at the same rate as wild-type animals, we examined the possibility that the rate of elimination could be slower, causing more bacteria to accumulate in the gut, by conducting defecation assays with young and old *parkin* mutants, as well as with wild-type controls. To measure the rate of defecation, cohorts of 40 animals per age and genotype were placed on fly food containing FD&C Blue Dye #1. After 24h allowing the blue food to reach steady state in the gut, animals were transferred to fresh blue food vials, and the number of blue fecal spots deposited on the walls of the vial was counted after 24h. Food vials were laid on their side, so that the climbing defects of *parkin* mutants would not affect the results of the experiment. We observed that young *parkin* mutants had significantly lower rates of defecation compared to wild-type controls (Supplementary Fig. S1). Older flies showed no difference in defecation rate, and, together with cell-type specific *parkin* RNAi experiments (see below), these results suggested elimination from the gut is unlikely to be the sole contributor to the elevated microbial load in *parkin* mutants.

### *Parkin* is required in gut enterocytes to maintain microbial load homeostasis

To determine which specific cell types required *parkin* activity to maintain gut microbial homeostasis, we characterized a *parkin* RNAi line and confirmed that ubiquitous *parkin* knockdown using this line led to a decrease in *parkin* RNA expression, muscle degeneration reflective of *parkin* loss of function, as well as the increase in gut microbial load (Fig. 2a-g). We then examined the role of tissues implicated in *parkin* function (the nervous system, muscle), as well as specific cell types within the gut for a role in the gut microbial phenotype. Knockdown of *parkin* in gut enterocytes (*NP1-GAL4* driver) resulted in the increased microbial load (Fig. 2h), whereas we observed no change in microbial load upon *parkin* depletion in gut stem cells (*esg-GAL4* driver), neurons (*elav-GAL4* driver), or muscle (*24B-GAL4* driver) (Fig. 2i-k). These results suggest that *parkin* gene function is required in gut enterocytes to maintain microbial load within the wild-type range.

**Figure 2.**
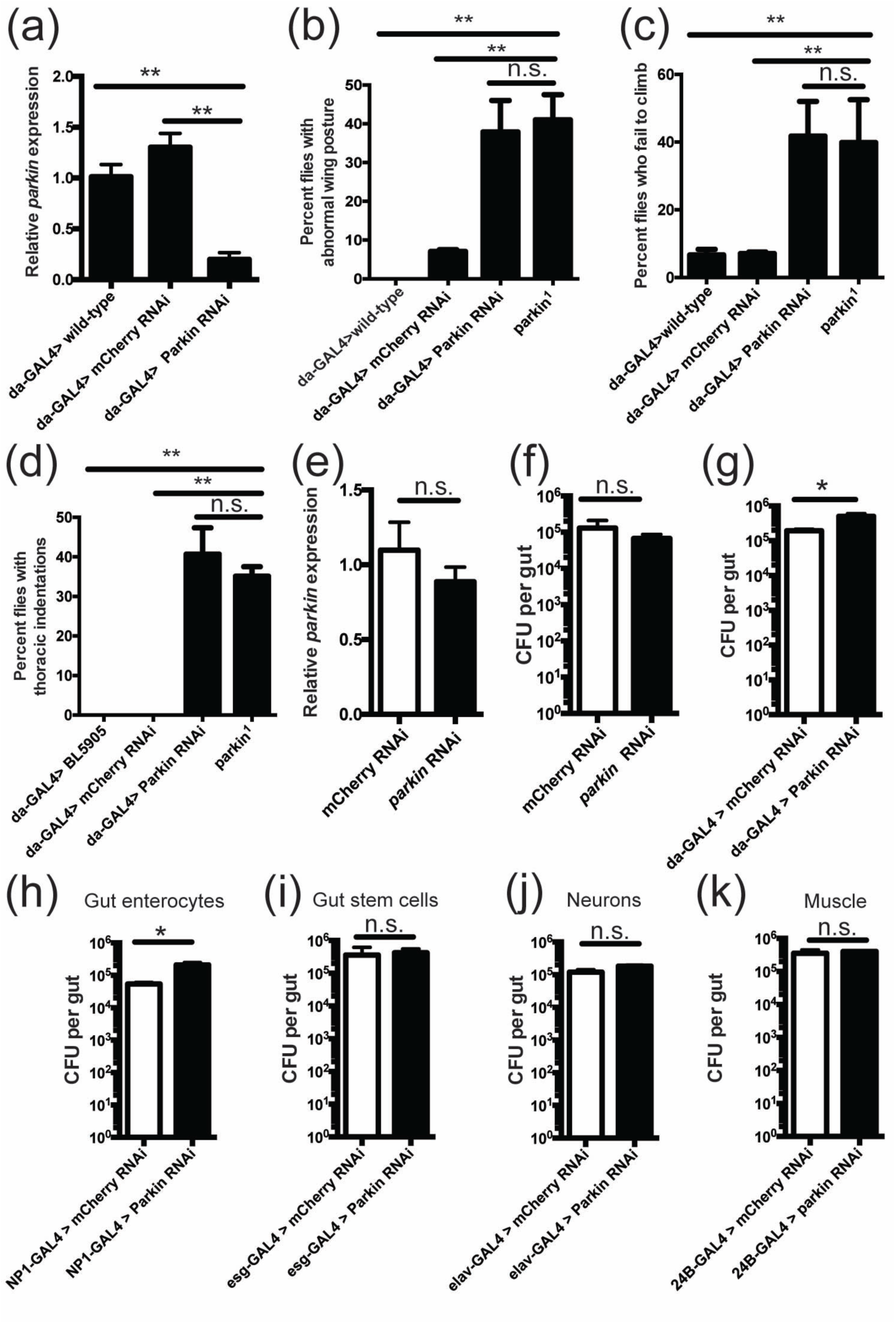
Parkin is required in gut enterocytes to maintain microbial homeostasis. **(A-D)** Validation of effective knockdown of *parkin* by in vivo expression of *siRNA* hairpin. (A) Real-time PCR for *parkin* in total RNA from whole 7d males expressing no hairpin, a hairpin against mCherry, or a hairpin against *parkin*. **p<0.01, ANOVA followed by Tukey’s post test. (B) Fraction of flies exhibiting abnormal wing posture among 7d males expressing no hairpin, a hairpin against mCherry, or the hairpin against *parkin*, compared to *parkin*^*1*^ flies. Flies were aged on standard food and the number of animals with held-up wings was counted. The experiment was conducted in biological triplicate using 55-60 flies per replicate. **p<0.01, n.s. – not significant, ANOVA followed by Tukey’s post test. (C) Climbing assay of 7d males expressing no hairpin, a hairpin against mCherry, or the hairpin against *parkin*, as well as *parkin*^*1*^ null flies. Flies were aged on standard food and placed in empty vials. The number of animals that climbed to the top of the vial 10s after tapping was recorded. Experiment was repeated in 3 independent biological replicates with 55-60 flies per replicate. **p<0.01, n.s. – not significant, ANOVA followed by Tukey’s post test. (D) Fraction of flies exhibiting thoracic indentations among 7d males expressing no hairpin, a hairpin against mCherry, or the hairpin against *parkin*, compared to *parkin*^*1*^ flies. Flies were aged on standard food and the number of animals with a collapsed thorax was counted. Experiment was repeated in 3 independent biological replicates with 55-60 flies per replicate. **p<0.01, n.s. – not significant, ANOVA followed by Tukey’s post test. **(E)** Real-time PCR for *parkin* in total RNA from whole 7d control or *parkin* RNAi males, in which the UAS-hairpin line was crossed to a wild-type line with no driver. n.s. – not significant, Student’s t-test. **(F)** Gut microbial load of 20d control or *parkin* RNAi males, in which the UAS-hairpin line was crossed to a wild-type line with no driver. n.s. – not significant, Student’s t-test. **(G)** Gut dissection followed by live colony counting in flies expressing mCherry or *parkin* RNAi ubiquitously. The gut dissection procedure was as in Fig 1A. The experiment was repeated in four independent biological replicates of six individual guts each per age and genotype. *p<0.05, Student’s t-test. **(H-K)** Microbial load in guts of 20d control or *parkin* RNAi males, in which knockdown was carried out selectively in (H) gut enterocytes, (I) gut stem cells, (J) neurons, or (K) muscle cells with indicated GAL4 drivers. Guts were dissected as in Fig 1A. The experiment was repeated in four independent biological replicates of six individual guts each per age and genotype. *p<0.05, n.s. – not significant, Student’s t-test.

### The gut microbiota impact *parkin* sensitivity to paraquat

The fly gut microbiota are beneficial for the host, promoting larval development under conditions of nutrient scarcity (Shin et al., 2011; Téfit and Leulier, 2017). We considered whether the increased microbial abundance in *parkin* flies may contribute to the *parkin* mutant phenotype. To assess this, we created germ-free animals by dechorionation of embryos followed by rearing on food supplemented with antibiotics (Guo et al., 2014; Ren et al., 2007). Flies mutant for *parkin* have a known increased sensitivity to oxidative toxins such as paraquat (Pesah et al., 2004). We assessed whether this phenotype was altered in germ-free animals, by subjecting germ-free and conventionally raised male flies to a paraquat sensitivity assay. Interestingly, we found that germ-free *parkin* flies survived longer on paraquat compared to conventional *parkin* animals (Fig. 3a). This finding suggests that the gut microbiota increase sensitivity of the *parkin* mutant to paraquat stress.

**Figure 3.**
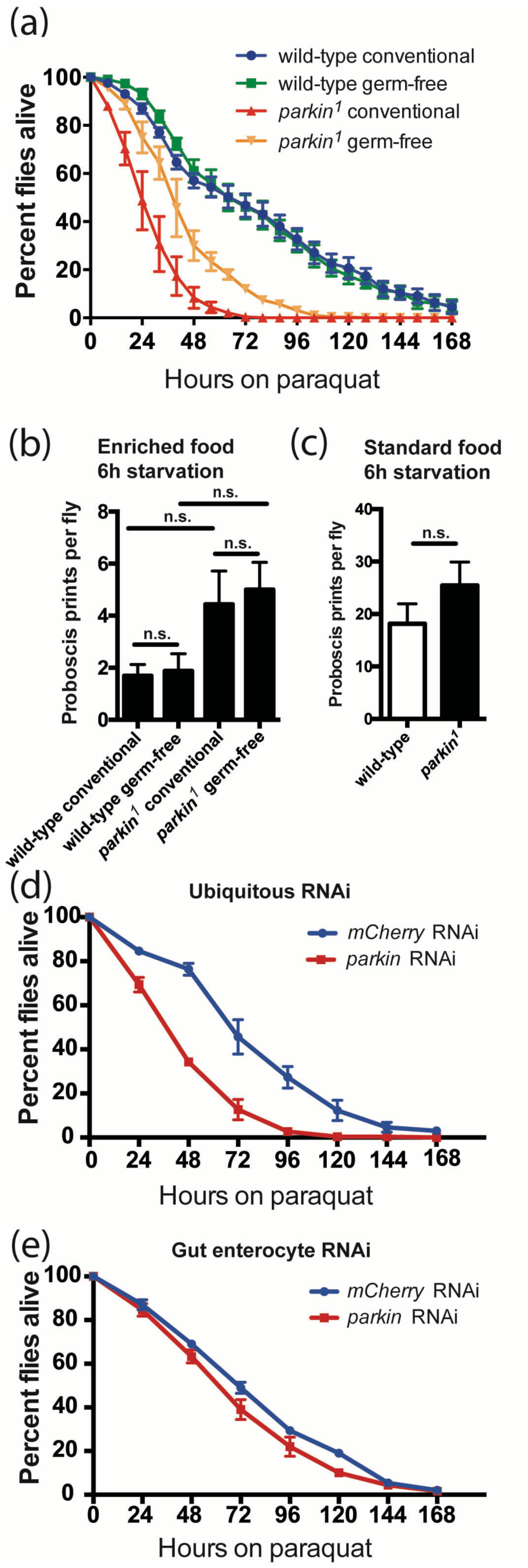
Absence of the gut microbiota affects paraquat sensitivity of *parkin* mutants. **(A)** Survival curve on 20 mM paraquat of 0-3d conventional or germ free wild-type and *parkin*^*1*^ mutant males. 100 animals per treatment and genotype were starved for 6h then placed on 10% sucrose-, 2.5% agar-food containing 20 mM paraquat. Survival was measured every 8h for 168h (over 7d). The experiment was repeated in three independent biological replicates. *parkin* conventional and germ-free animals had significantly different survival curves to one another and to their respective wild-type controls (p<0.0001, Log-Rank test). **.(B)** Proboscis print assay to measure the rate of feeding of conventionally reared and germ-free wild-type and *parkin*^*1*^ mutant males at ages 0-3d. Assay was carried out as in Fig 1 but with flies grown on food supplemented with 100 g/L yeast and starved for 6h prior to the assay. n.s. - not significant, ANOVA followed by Tukey post-test. **(C)** Proboscis print assay to measure the rate of feeding of control and *parkin*^*1*^ mutant males at ages 0-3d grown on standard fly food and starved for 6h prior to the assay. n.s. - not significant, Student’s t-test. **(D-E)** Paraquat sensitivity assays with 0-3d control or *parkin* RNAi males, in which knockdown was carried out ubiquitously (D, da-GAL4 driver) or selectively in gut enterocytes (D, NP1-GAL4 driver). Paraquat sensitivity assays were carried out as in Fig 3A. The experiment was repeated in three independent biological replicates of 100 individual animals each per experimental group. (D) ***p<0.0001, log-rank test. (E) Not significant, log-rank test.

We confirmed that improved paraquat resistance of germ-free flies was not due to the animals eating less and thus ingesting less of the toxin, as proboscis print assays showed no difference in the rate of feeding between germ-free and conventional *parkin* males (Fig. 3B). The proboscis print assay showed no significant difference in feeding between *parkin* and wild-type males unlike the previous assay that showed parkin flies eat less (see Fig. 1d). Proboscis print assays on wild-type and *parkin* males reared on standard food and treated with starvation caused no difference in feeding rate analogous to the assay with males from germ-free lines (Fig. 3c), leading us to conclude that *parkin* and wild-type flies eat equally in response to starvation.

We further investigated whether *parkin* knockdown in the gut selectively affects paraquat sensitivity, or alternatively, if paraquat sensitivity is a non-gut phenotype that is affected by the presence of the gut microbiota. To examine this, we used conditional *parkin* RNAi followed by paraquat sensitivity assays. Ubiquitous RNAi of *parkin* phenocopied the increased toxin sensitivity of the *parkin* mutant (Fig. 3d). Intriguingly, *parkin* RNAi knockdown selectively in gut enterocytes did not cause a significant change in paraquat sensitivity (Fig 3E). Taken together with a recent study suggesting that increased paraquat sensitivity in *parkin* mutants may be due to *parkin* loss of function in muscle and brain (de Oliveira Souza et al., 2017), these results indicate that paraquat sensitivity is not a gut-specific effect but that altering the gut microbiota can influence non-gut animal characteristics, namely sensitivity to toxins.

### The gut microbiota are altered in composition in aged parkin mutants

Given the impact of *parkin* gene function on gut microbial abundance, we determined whether there were alterations in the composition of microbes in the *parkin* gut. To define the microbial types, we sequenced 16S rDNA V1-V2 variable region amplicons using DNA extracted from dissected guts of 7d and 20d wild-type and *parkin* males. For the young timepoint, we chose 7d rather than 3d due to the very low microbial abundance in 3d guts. Sequences were clustered into Operational Taxonomic Units (OTUs) by aligning against “seed” sequences from the Greengenes database (Caporaso et al., 2010), or if clustering with Greengenes failed, by aligning against each other (open-reference OTU picking). The taxonomic identity of each OTU was assigned using the RDP classifier(Wang et al., 2007). We found no significant difference in α-diversity between *parkin* and wild-type microbiomes using several diversity metrics (Supplementary Table S1). Weighted UniFrac showed no difference at 7d in microbial composition between *parkin* null and control males (Fig 4A). At 20d, however, the composition of the gut microbiota of *parkin* nulls and wild-type flies diverged from each other and from the microbiome of 7d males (Fig. 4a). These data indicate that aged parkin mutants not only have a higher gut bacterial load, but also an altered gut genera composition compared to normal animals.

**Figure 4.**
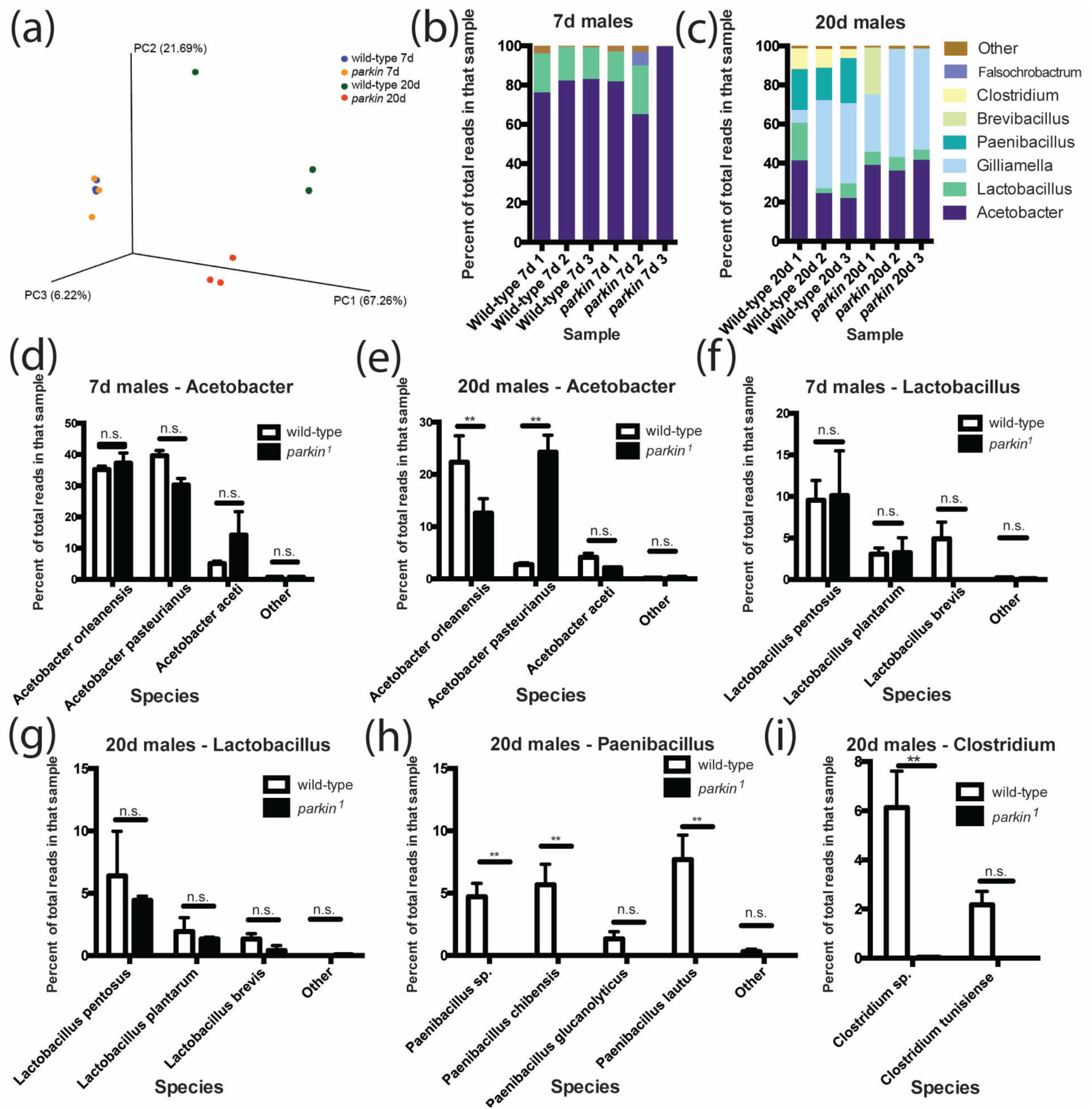
*parkin* loss of function affects gut microbial composition. 16S rDNA amplicon sequencing of guts from 7d and 20d wild-type and *parkin* males. **(A)** Principle coordinate analysis shows similar microbial composition at age 7d, but at age 20d compositions of the gut microbiota of wild-type and *parkin*^*1*^ mutant diverge. **(B-C)** Most common genera (defined as more than 5% of total reads in at least one sample) in (B) 7d male guts and (C) 20d male guts. **(D-I)** Relative abundance (measured as percentage of total reads in that sample) of *Acetobacter* species detected in (D) 7d males and (E) 20d males, *Lactobacillus* species detected in (F) 7d males and (G) 20d males, *Paenibacillus* species detected in (H) 20d males, and *Clostridium* species detected in (I) 20d males. **p<0.01, n.s. – not significant, Student’s t-test with Holm-Sidak correction for multiple testing.

We defined the variation underlying the divergent microbiome of aged *parkin* animals by analyzing the most abundant gut genera, defined as comprising at least 5% of the total reads in any one sample. These data showed that 20d *parkin* mutants have a decreased relative abundance of *Paenibacillus* and *Clostridium* reads (Fig. 4c). To interrogate differences at the species level, representative sequences from each OTU were fetched and batch-aligned to the BLAST 16S rRNA sequence database using nucleotide BLAST. The top hit with more than 99% identity to a sequence from an identified species in the database, defined the species identity and was assigned to the OTU (see Supplementary Figures S2-S4 for representative alignments). Species-level analysis revealed a switch of the dominant *Acetobacter* species from *A. orleanensis* to *A. pasteurianus* in 20d *parkin* males (Fig. 4e).

## DISCUSSION

In this study we examined the relationship between microbes in the gut and *parkin* gene function. We discovered a five-fold increase of microbial load in the guts of aged *parkin* flies compared to wild-type controls. *In vivo* RNAi of *parkin* in gut enterocytes revealed that *parkin* gene function in the gut specifically impacts microbial load. Paraquat sensitivity assays with germ-free flies showed a beneficial effect on paraquat sensitivity in germ-free *parkin* animals compared to conventionally reared controls. Using 16S rDNA sequencing, we assessed the effect of the *parkin* mutation on gut microbial composition and observed an altered bacterial genera and species abundance in aged *parkin* flies.

Unexpectedly, the increase compared to controls of live microbes in the guts of 20d *parkin* flies was not also observed in *pink1* flies, even though Pink1 and Parkin share many age-associated adult-onset phenotypes, and regulate mitophagy and mitochondrial fission/fusion as parts of the same pathway (Pickrell and Youle, 2015). In mammals, Parkin has been shown to ubiquitinate and activate NEMO, a member of the NF-κB pathway, in a manner that is independent of Pink1 function (Müller-Rischart et al., 2013). Parkin also mediates ubiquitination of intracellular pathogens; whether Pink1 is required for this activity is not known (Manzanillo et al., 2013). Taken together, these observations suggest that Parkin has roles that are independent of Pink1 gene function; regulation of microbial homeostasis may be one such function.

Our data suggest that *parkin* gene function impacts gut microbial load and abundance. There are a number of ways in which an increase in microbial load may be linked to a change in microbial composition. The increase may lead to a spike in inflammation and oxidative stress, rendering the gut inhospitable for some taxa that otherwise would be present. It is also possible that *parkin* loss of function causes a decrease in relative abundance of some microbes that would normally limit proliferation of other taxa, leading to overgrowth of the remaining taxa.

It is unlikely that the effects of *parkin* loss of function on the gut microbiota are secondary effects of the known function of *parkin* to disrupt mitochondrial homeostasis, since *pink1* mutants have similar effects on the mitochondria but not microbial load. Although we cannot fully rule out an effect on microbiota due to a change in defecation rate, we speculate that Parkin may regulate gut microbial homeostasis via interactions with *Drosophila* innate immunity pathways. Two immunity pathways are known to regulate microbes in the fly gut: the Dual oxidase (Duox) and Imd pathways (Broderick and Lemaitre, 2012). Duox, a member of the NADPH oxidase family, produces reactive oxygen species that restrict bacterial viability(Kim and Lee, 2014). The enzyme activity is known to be upregulated by bacterial-derived uracil(Lee et al., 2015). To our knowledge, no link between Parkin activity and Duox is known at present. Alternatively, the Imd pathway is the *Drosophila* analog of the mammalian NF-κB pathway(Myllymäki et al., 2014). In the fly, the pathway promotes transcription and ultimately secretion of antimicrobial peptides (AMPs) in response to DAP-type peptidoglycan, a component of bacterial cell walls (Myllymäki et al., 2014). Interestingly, Parkin in mammals ubiquitinates a member of the NF-κB pathway, NEMO (Müller-Rischart et al., 2013), which is essential for NF-κB pathway activation. This activity is independent of Pink1 (Müller-Rischart et al., 2013). The *Drosophila* NEMO homolog, IKK-γ, also plays a role in activation of the Imd pathway (Ertürk-Hasdemir et al., 2009; Rutschmann et al., 2000). In mice, conditional ablation of NEMO leads to impaired AMP secretion, intestinal epithelial cell apoptosis, and translocation of bacteria into the intestinal mucosa (Nenci et al., 2007).

A surprising result is that the presence of a gut microbiome is detrimental to *parkin* mutants exposed to paraquat. Given the improved toxin resistance of germ-free *parkin* flies, metabolism of paraquat by microbes found in the *parkin* gut may increase paraquat toxicity. Many bacteria have been shown to be able to use paraquat as an electron carrier in the redox cycle, generating reactive oxygen species (ROS) (Haley, 1979). ROS generated by gut bacteria through redox cycling would not only be toxic in themselves, but also increase gut permeability, allowing even more toxic paraquat to be taken up by the fly. Paraquat can also be used as a coenzyme by bacteria in the reduction of sulfate, thiosulfate, hydroxylamine, nitrate, among other compounds (Haley, 1979). It is possible that paraquat could mediate increased secretion of a gut bacterial metabolite which in turn is toxic to the host.

Our results suggest Parkin plays a before-undocumented role in regulation of gut microbial homeostasis, and conversely, that the gut microbiota impact parkinsonism as modeled in the fly. This study deepens our understanding of the *parkin* mutant phenotype and sets a foundation for further studies on the importance of the gut microbiota to parkinsonism in mammals.

## Acknowledgements

We thank members of the Bonini laboratory for critical reading and helpful comments. We thank Karl Kumbier (UCB Statistics Department) and Jian-Hua Mao for reviewing our statistical analyses. This work was supported by funding from NIH grants R21-NS088370, R35-NS097275, and a Glenn Award for Research in Biological Mechanisms of Aging (to N.M.B.). Additional support was provided by Lawrence Berkeley National Laboratory (LBNL) Directed Research and Development (LDRD) program funding under the Microbes to Biomes (M2B) initiative (K.H.W. and S.E.C.). LBNL is a multi-program national laboratory operated by the University of California for the DOE under Contract DE AC02-05CH11231.

## Author Contributions

V.F. and N.M.B. conceived and designed experiments. V.F. performed experiments and analyzed data. K.H.W. prepared and sequenced libraries. S.E.C. provided input, experimental advice and equipment. V.F. and N.M.B. wrote the manuscript with input from S.E.C.

## METHODS

### Fly lines

Flies were grown in standard cornmeal-molasses-agar medium at 25°C. *parkin*^*1*^ (*w*\*;; *P[EP]park1/TM3, Sb1 Ser1*, FlyBase ID: FBst0034747) and *parkin* RNAi *(y1 sc*\* *v1; P[TRiP*.*HMS01800]attP2/TM3, Sb1*, FlyBase ID: FBst0038333) flies were obtained from the Bloomington Stock Center. NP1-GAL4 flies were obtained from Sara Cherry (University of Pennsylvania) and *pink (pink1*^*B9*^*/FM6)* flies were obtained from Jongkyeong Chung (Seoul National University). The *parkin*^*1*^ and *pink1*^*B9*^ alleles were backcrossed into a homogenous wild-type background (*w*^*1118*^, FlyBaseID: FBst0005905) for five generations. DJ1 DKO (*w*^*1118*^;*DJ-1α*^*Δ72*^; *DJ-1β*^*Δ93*^*/TM6,Tb*) flies are described (Meulener et al., 2005).

### Gut dissection and CFU counting

Animals used in different replicates were collected from different bottles and aged in different vials. All animals were collected as virgins and aged on standard cornmeal-molasses-agar medium at a density of 20 flies per vial. Fly density has been shown to impact the relative abundances of *Acetobacter* and *Lactobacillus* (Wong et al., 2015). Animals were transferred to fresh food vials every other day. *parkin* and wild-type animals were aged in the same vials. For all experiments, controls and experimentals were aged on the same batch of food and transferred on the same day. For the tissue specific expression experiments, the driver line was compared to the RNAi line using the same food and under the same conditions, as different driver lines represent different background.

For gut dissection, flies were anesthetized, washed 1X in 1mL 10% bleach, 1X in 1mL 100% ethanol, and finally rinsed 3X in 1 mL sterile PBS (Sigma-Aldrich). 200 µL of the final rinse were spread on an MRS-agar plate (BD Diagnostic Systems) as a control for the efficacy of the wash. Each gut was dissected in a drop of PBS on a sterile microscope slide and placed in 200 µL PBS. The gut was homogenized by bead-beating with 1mm tissue-disruption beads (Research Products International) for 30s at maximum speed. 10-, 100-, and 1000-fold dilutions of the homogenate were spread on MRS-agar plates. The fly gut is aerobic, allowing culture of bacteria with most standard media and conditions, including the microbes defined here (Guo et al., 2014; He et al., 2007). All plates were incubated at 30°C for 48h. Bacterial colonies were counted and multiplied by the dilution factor to calculate the number of Colony Forming Units (CFU) per gut.

### Proboscis prints, blue dye, and defecation assays

Proboscis print assays were modified from Edgecomb et al. (1994). Clean microscope slides (Fisher) were briefly dipped in 10% sucrose 1% gelatin and left to dry at room temperature in a covered area for 3-4h. Flies were anesthetized and placed in individual wells of a 96-well plate. Groups of ten flies were arranged in two columns of five wells. Each group was covered by a strip of wax paper and a gelatinated microscope slide. Flies were incubated for 30 min at room temperature to recover from the anesthesia, during which time the outline of each well was traced on the slide using a thin permanent marker. At the end of the incubation period, the strip of wax paper was swiftly removed allowing contact between the fly and the sweet gelatin coat. Plates were inverted, allowing the flies to walk on top of the slide for 20 min at room temperature. The number of prints left on each slide was counted using Differential Interference Contrast (DIC) microscopy.

For the blue dye assays, flies were fed for 48h on standard food supplemented with 2.5% w/v FD&C Blue Dye #1 (SPS Alfachem). Five guts per age and genotype were dissected, homogenized and the absorbance of the sample at 630nm was measured with a spectrophotometer.

For the defecation assays, cohorts of 40 animals per age and genotype were tested using ten flies per vial on fly food containing 2.5% w/v FD&C Blue Dye. Animals were left on the dye for 24h. Flies were transferred to fresh blue food vials and the number of blue fecal spots deposited on the walls of the vials was counted after 24h.

### Germ-free flies

The germ-free fly protocol was adapted from previously described techniques.(Guo et al., 2014; Koyle et al., 2016; Ma et al., 2015) Standard cornmeal-molasses-agar fly food was autoclaved and upon cooling supplemented with yeast extract (Fisher) to 100 g/L. An antibiotic cocktail of kanamycin (1mM; Fisher), ampicillin (650 µM; MediaTech), and doxycycline (650 µM; Sigma-Aldrich) was added to the food as previously described(Ren et al., 2007). Food was dispensed in empty fly bottles at 50 mL per bottle in a laminar flow cabinet and left to solidify. A 12h collection of fly embryos was rinsed in 100% ethanol to cleanse and sterilize any leftover agar from collection plates, dechorionated in 10% bleach for 2 min, and immediately rinsed 3X in sterile PBS. Embryos were placed on the prepared fly food and overlaid with sterile glycerol. Germ-free fly lines were maintained on sterile food for up to 3-4 generations using a laminar flow cabinet. Flies were monitored for bacterial contamination by homogenizing larvae and testing for bacterial growth on MRS-agar plates.

### Paraquat sensitivity assays

Flies were transferred to empty vials at 20 flies per vial (Genesee Scientific), starved for 6h, then transferred to vials containing 2.5% agar (LabScientific), 10% sucrose (Sigma-Aldrich), 25mM Paraquat (MP Biomedicals). Vials were incubated at 25°C and the number of dead flies in each vial was counted every 8h until all flies were dead or until 168 hr (7d) had passed.

### 16S rDNA sequencing

Animals from different replicates were collected from different bottles and aged in different vials to ensure replicates were biologically independent. Flies were aged at a density of 20 flies per vial and transferred to fresh food vials every other day. Wild-type and *parkin* flies were aged on the same batch of food and transferred at the same time. All twenty flies from a vial were used for each biological replicate. Twenty guts per sample were dissected as described above and subjected to DNA extraction using the PSP Spin Stool DNA Purification Kit (Stratec Biomedical). PCR of the V1-V2 variable regions was performed using the 27F – 338R primer pair (27F: 5′-AGAGTTTGATCMTGGCTCAG-3′; 338R: 5′-TGCTGCCTCCCGTAGGAGT-3′) with the following program: 94°C for 4 min, 94°C for 30s, 58°C for 30s, 72°C for 40s, 30 total amplification cycles, 72°C for 10 min, then hold 4°C. Three PCR reactions were pooled and the PCR product was purified using the Agencourt AMPure XP PCR purification kit (Beckman Coulter) and sequenced using MiSeq (Illumina).

Sequencing analysis was carried out using the QIIME suite(Caporaso et al., 2010). Paired reads were joined and quality filtered using a Phred score cutoff of 20. OTUs were picked using an open-reference OTU picking algorithm with the Uclust alignment method and 99% identity. OTUs with less than 10 reads were removed from the analysis. The most abundant sequence was selected as a representative sequence for each OTU and used to assign a taxonomic classification for each OTU using the RDP classifier version 2.12(Wang et al., 2007). The resulting OTUs and their taxonomy were compiled in a QIIME OTU table.

### Climbing assays, thoracic indentations, and abnormal wing posture scoring

Flies were raised and aged on standard cornmeal molasses agar food vials at a density 20 flies per vial. Number of flies with abnormal wing posture was scored on anaesthetized animals in the vial. For climbing assays, flies were flip-transferred in empty vials (Genesee Scientific) with a line marking a distance 8 cm above the bottom of the vial, near the top. Vials were tapped and the number of animals that crossed the mark 10s after tapping was recorded. The presence or absence of thoracic indents was scored on anesthetized animals on a fly pad. Experiments were repeated in 3 independent biological replicates with 55-60 flies per replicate.

### Real-time quantitative PCR

Total RNA from crushed whole males was purified using the Trizol reagent (Ambion) following the reagent manual. The RNA was DNase treated using TURBO DNase (Ambion) according to the kit instructions. After DNase treatment, the RNA was Trizol purified again. Reverse transcription was carried out using the High-Capacity cDNA Reverse Transcription Kit (Applied Biosystems) according to the kit manual. Real time PCR was carried out using the Fast SYBR Green Mastermix (Applied Biosystems) following the kit instructions. Primers used for RT PCR had the following sequences: *parkin* F: 5’-CGGATGTGAGTGATACCGTGT-3’; *parkin* R: 5’-ATAAACTGACGCTCGCCCAA-3’.

### Statistics

Statistical analyses pertaining to the processing of 16S rDNA sequencing results were carried out using QIIME’s built-in functions(Caporaso et al., 2010). All other statistical tests were performed using GraphPad Prism (GraphPad Software, La Jolla, CA). For treatment and mutant analyses, we used the Analysis of Variance (ANOVA) test to determine differences between three or more means. If significance was detected, Tukey’s post-test was used to identify those values that were significantly different.

---

**Supplementary Table 1.**

|  | Simpson index | Shannon index | Chao1 | PD whole tree |
| --- | --- | --- | --- | --- |
| Wild-type 7d | 0.71±0.02 | 2.36±0.12 | 327.78±36.57 | 10.81±1.06 |
| *parkin* 7d | 0.72±0.03 | 2.31±0.29 | 299.20±71.22 | 10.81±1.06 |
| p-value | 0.69 | 0.91 | 0.74 | 0.82 |
| Wild-type 20d | 0.80±0.03 | 3.18±0.13 | 379.29±29.14 | 11.74±0.61 |
| *parkin* 20d | 0.69±0.06 | 2.34±0.33 | 327.72±8.44 | 10.69±0.54 |
| p-value | 0.24 | 0.11 | 0.21 | 0.27 |

Multiple metrics were used to best capture and compare the biodiversity that exists between the different gut bacterial communities. Diversity indices of wild-type and *parkin* 7d and 20d gut microbiomes using measures available from QIIME (Caporaso et al., 2010). The Simpson index is used to measure species richness (number of different species) and the Shannon index is used to measure species evenness (relative abundance of species). The Chao1 metric is used to analyze data sets with low-abundance classes. PD whole tree assesses phylogenetic diversity. Values are means ± SEM, n=3. *p*-values were calculated using Student's *t*-test comparing wild-type and *parkin* values at the specified age. The *p*-values are all >.05 demonstrating no significant differences between *parkin* and wild-type flies.

**Supplementary Figure 1:**
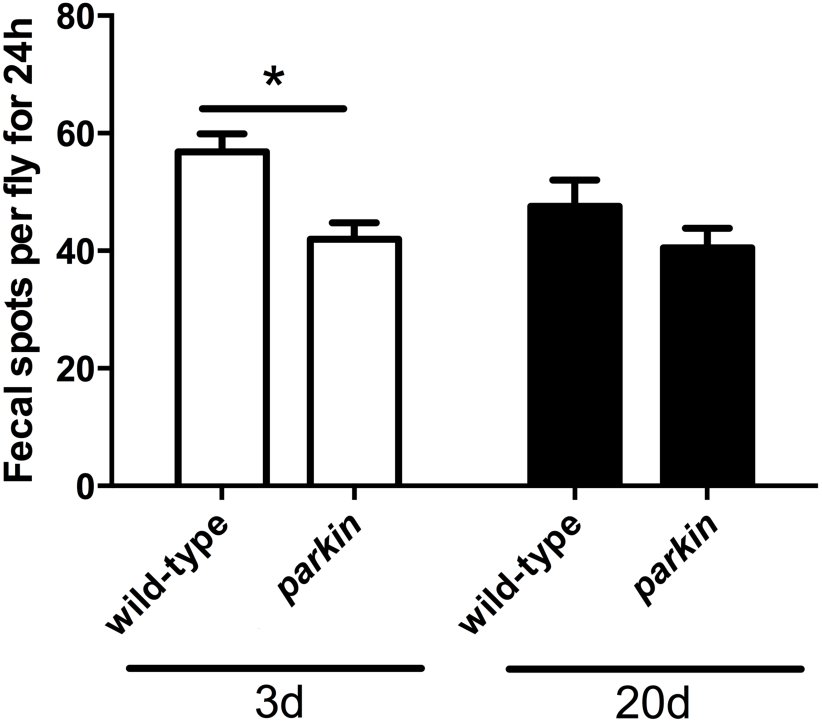
Decreased defecation rate in young *parkin* male guts. Cohorts of 40 males were incubated on fly food containing FD&C Blue Dye #1 for 24h. Flies were transferred to fresh blue food vials, and after another 24h incubation period, the number of blue fecal spots on the walls of the vials were counted. The experiment was repeated in four independent biological replicates. *p<0.05, ANOVA with Tukey’s post test.

**Supplementary Figure 2:**
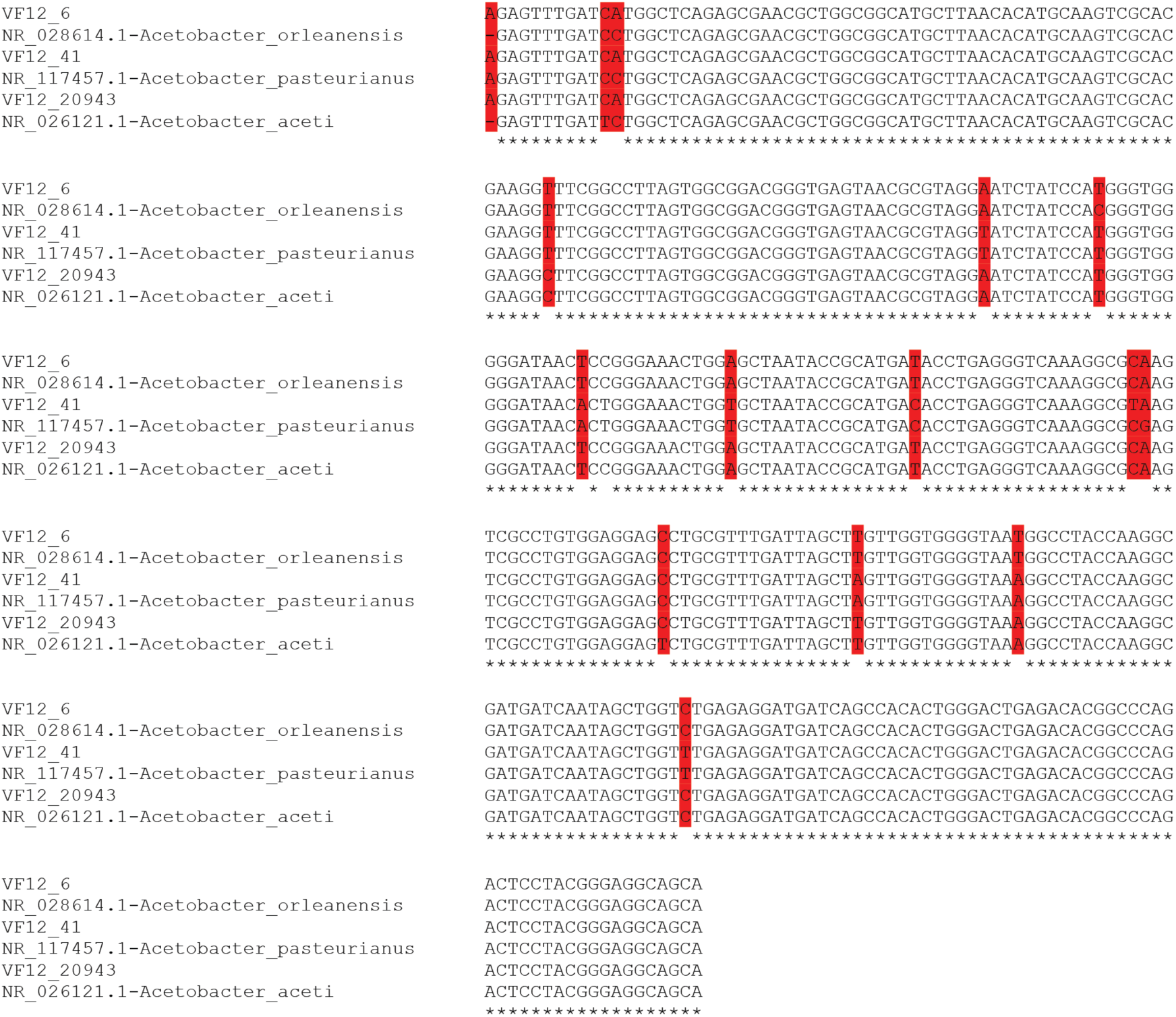
Alignment between sequenced *Acetobacter* 16S rDNA amplicons and best matches from BLAST search. Reads were fetched from the set of representative sequences for each OTU and BLAST searched against the NCBI 16S rDNA sequence database. Three pairs of reads and their BLAST top hit were aligned using Clustal Omega. Mismatching nucleotides that can be used to differentiate between species are highlighted in red. VF12_6 was identified as *Acetobacter orleanensis*. VF12_41 was identified as Acetobacter *pasteurianus*. VF12_20943 was identified as *Acetobacter aceti*.

**Supplementary Figure 3:**
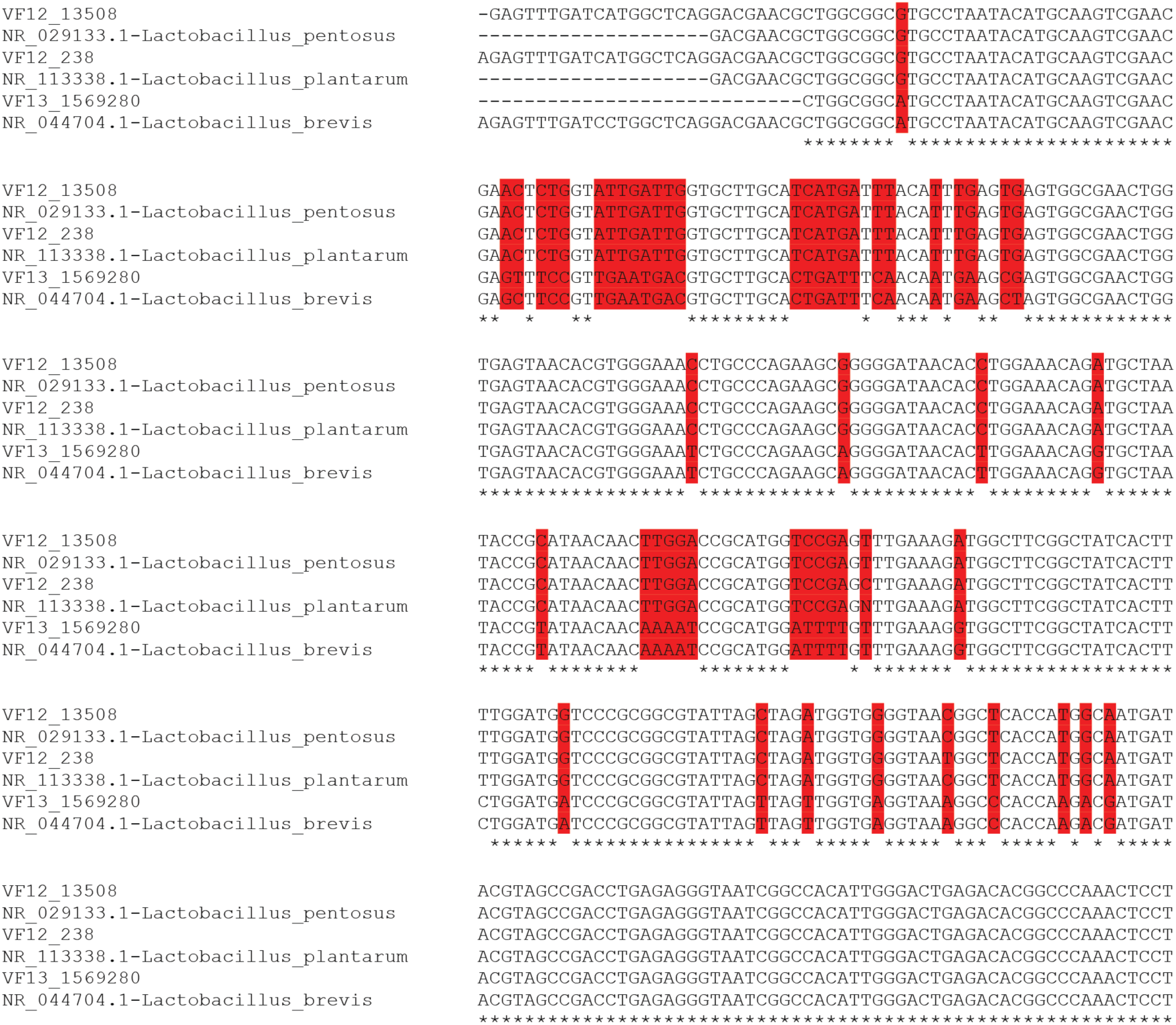
Alignment between sequenced *Lactobacillus* 16S rDNA amplicons and best matches from BLAST search. Reads were fetched from the set of representative sequences for each OTU and BLAST searched against the NCBI 16S rDNA sequence database. Three pairs of reads and their BLAST top hit were aligned using Clustal Omega. Mismatching nucleotides that can be used to differentiate between species are highlighted in red. VF12-13508 was identified as *Lactobacillus pentosus*. VF12_238 was identified as *Lactobacillus plantarum*. VF13_1569280 was identified as *Lactobacillus brevis*.

**Supplementary Figure 4:**
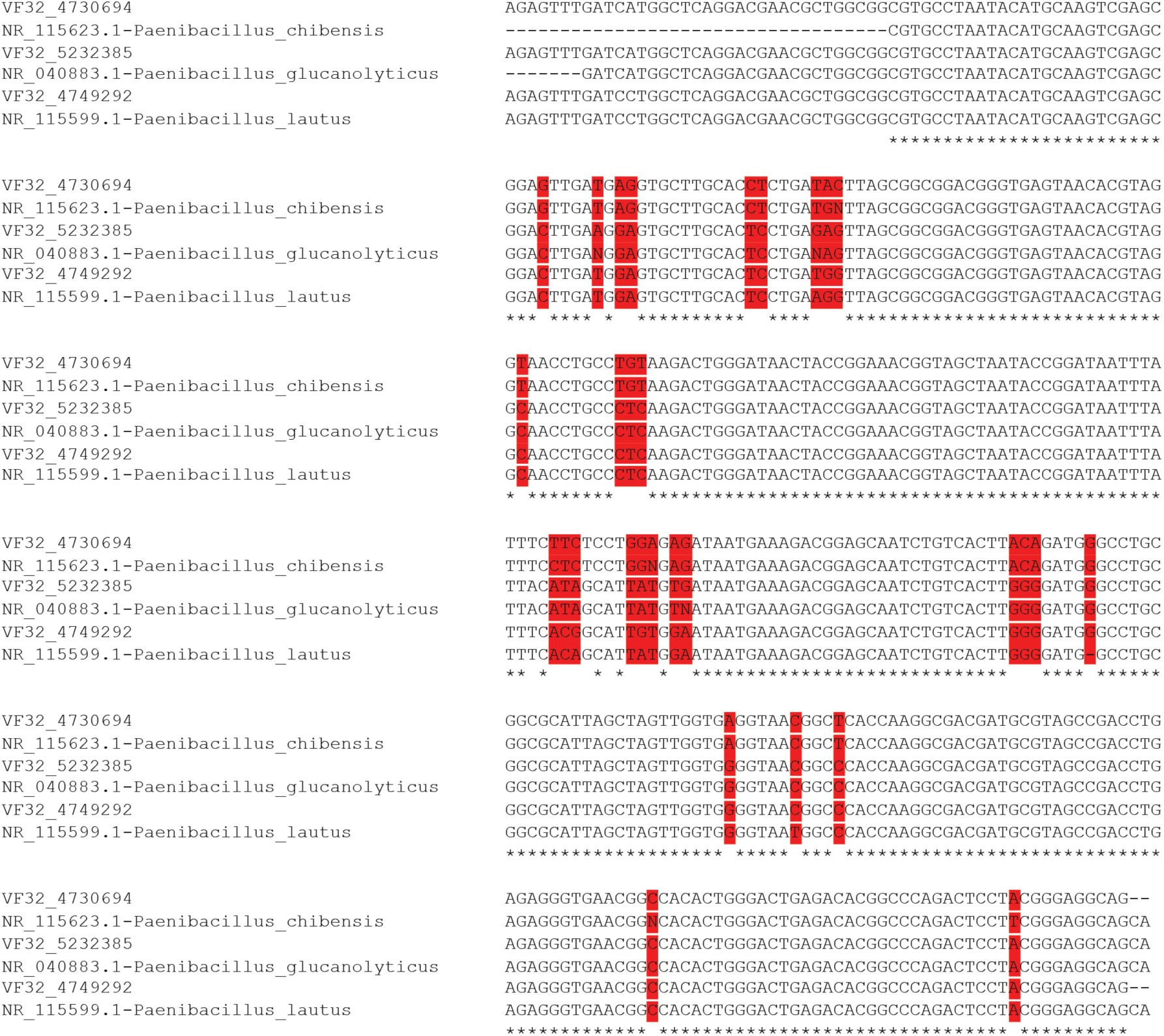
Alignment between sequenced *Paenibacillus* 16S rDNA amplicons and best matches from BLAST search. Reads were fetched from the set of representative sequences for each OTU and BLAST searched against the NCBI 16S rDNA sequence database. Three pairs of reads and their BLAST top hit were aligned using Clustal Omega. Mismatching nucleotides that can be used to differentiate between species are highlighted in red. VF32_4730694 was identified as *Paenibacillus chibensis*. VF32_5232385 was identified as *Paenibacillus glucanolyticus*. VF32_4749292 was identified as *Paenibacillus lautus*.

## REFERENCES

Arumugam, M., Raes, J., Pelletier, E., Le Paslier, D., Yamada, T., Mende, D.R., Fernandes, G.R., Tap, J., Bruls, T., Batto, J.M., et al. (2011). Enterotypes of the human gut microbiome. Nature 473, 174–180.

Bier, E. (2005). *Drosophila*, the golden bug, emerges as a tool for human genetics. Nat Rev Genet 6, 9–23.

Bravo, J.A., Forsythe, P., Chew, M.V., Escaravage, E., Savignac, H.M., Dinan, T.G., Bienenstock, J., and Cryan, J.F. (2011). Ingestion of *Lactobacillus* strain regulates emotional behavior and central GABA receptor expression in a mouse via the vagus nerve. Proc Natl Acad Sci U S A 108, 16050–16055.

Broderick, N.A., Buchon, N., and Lemaitre, B. (2014). Microbiota-induced changes in *Drosophila melanogaster* host gene expression and gut morphology. MBio 5, e01117–01114.

Broderick, N.A., and Lemaitre, B. (2012). Gut-associated microbes of *Drosophila melanogaster*. Gut Microbes 3, 307–321.

Canet-Avilés, R.M., Wilson, M.A., Miller, D.W., Ahmad, R., McLendon, C., Bandyopadhyay, S., Baptista, M.J., Ringe, D., Petsko, G.A., and Cookson, M.R. (2004). The Parkinson’s disease protein DJ-1 is neuroprotective due to cysteine-sulfinic acid-driven mitochondrial localization. Proc Natl Acad Sci U S A 101, 9103–9108.

Caporaso, J.G., Kuczynski, J., Stombaugh, J., Bittinger, K., Bushman, F.D., Costello, E.K., Fierer, N., Pena, A.G., Goodrich, J.K., Gordon, J.I., et al. (2010). QIIME allows analysis of high-throughput community sequencing data. Nat Methods 7, 335–336.

de Oliveira Souza, A., Couto-Lima, C.A., Rosa Machado, M.C., Espreafico, E.M., Pinheiro Ramos, R.G., and Alberici, L.C. (2017). Protective action of Omega-3 on paraquat intoxication in *Drosophila melanogaster*. J Toxicol Environ Health A 80, 1050–1063.

Deng, H., Dodson, M.W., Huang, H., and Guo, M. (2008). The Parkinson’s disease genes pink1 and parkin promote mitochondrial fission and/or inhibit fusion in *Drosophila*. Proc Natl Acad Sci U S A 105, 14503–14508.

Eckburg, P.B., Bik, E.M., Bernstein, C.N., Purdom, E., Dethlefsen, L., Sargent, M., Gill, S.R., Nelson, K.E., and Relman, D.A. (2005). Diversity of the human intestinal microbial flora. Science 308, 1635–1638.

Edgecomb, R.S., Harth, C.E., and Schneiderman, A.M. (1994). Regulation of feeding behavior in adult *Drosophila melanogaster* varies with feeding regime and nutritional state. J Exp Biol 197, 215–235.

Ertürk-Hasdemir, D., Broemer, M., Leulier, F., Lane, W.S., Paquette, N., Hwang, D., Kim, C.H., Stöven, S., Meier, P., and Silverman, N. (2009). Two roles for the *Drosophila* IKK complex in the activation of Relish and the induction of antimicrobial peptide genes. Proc Natl Acad Sci U S A 106, 9779–9784.

Fink, C., Staubach, F., Kuenzel, S., Baines, J.F., and Roeder, T. (2013). Noninvasive analysis of microbiome dynamics in the fruit fly *Drosophila melanogaster*. Appl Environ Microbiol 79, 6984–6988.

Greene, J.C., Whitworth, A.J., Kuo, I., Andrews, L.A., Feany, M.B., and Pallanck, L.J. (2003). Mitochondrial pathology and apoptotic muscle degeneration in *Drosophila parkin* mutants. Proc Natl Acad Sci U S A 100, 4078–4083.

Guo, L., Karpac, J., Tran, S.L., and Jasper, H. (2014). PGRP-SC2 promotes gut immune homeostasis to limit commensal dysbiosis and extend lifespan. Cell 156, 109–122.

Haley, T.J. (1979). Review of the toxicology of paraquat (1,1’-dimethyl-4,4’-bipyridinium chloride). Clin Toxicol 14, 1–46.

Hayashi, T., Ishimori, C., Takahashi-Niki, K., Taira, T., Kim, Y.C., Maita, H., Maita, C., Ariga, H., and Iguchi-Ariga, S.M. (2009). DJ-1 binds to mitochondrial complex I and maintains its activity. Biochem Biophys Res Commun 390, 667–672.

He, Z., Kisla, D., Zhang, L., Yuan, C., Green-Church, K.B., and Yousef, A.E. (2007). Isolation and identification of a *Paenibacillus polymyxa* strain that coproduces a novel lantibiotic and polymyxin. Appl Environ Microbiol 73, 168–178.

Hsiao, E.Y., McBride, S.W., Hsien, S., Sharon, G., Hyde, E.R., McCue, T., Codelli, J.A., Chow, J., Reisman, S.E., Petrosino, J.F., et al. (2013). Microbiota modulate behavioral and physiological abnormalities associated with neurodevelopmental disorders. Cell 155, 1451–1463.

Jin, S.M., Lazarou, M., Wang, C., Kane, L.A., Narendra, D.P., and Youle, R.J. (2010). Mitochondrial membrane potential regulates PINK1 import and proteolytic destabilization by PARL. J Cell Biol 191, 933–942.

Kane, L.A., Lazarou, M., Fogel, A.I., Li, Y., Yamano, K., Sarraf, S.A., Banerjee, S., and Youle, R.J. (2014). PINK1 phosphorylates ubiquitin to activate Parkin E3 ubiquitin ligase activity. J Cell Biol 205, 143–153.

Kazlauskaite, A., Kondapalli, C., Gourlay, R., Campbell, D.G., Ritorto, M.S., Hofmann, K., Alessi, D.R., Knebel, A., Trost, M., and Muqit, M.M. (2014). Parkin is activated by PINK1-dependent phosphorylation of ubiquitin at Ser65. Biochem J 460, 127–139.

Kim, S.H., and Lee, W.J. (2014). Role of DUOX in gut inflammation: lessons from *Drosophila* model of gut-microbiota interactions. Front Cell Infect Microbiol 3, 116.

Kondapalli, C., Kazlauskaite, A., Zhang, N., Woodroof, H.I., Campbell, D.G., Gourlay, R., Burchell, L., Walden, H., Macartney, T.J., Deak, M., et al. (2012). PINK1 is activated by mitochondrial membrane potential depolarization and stimulates Parkin E3 ligase activity by phosphorylating Serine 65. Open Biol 2, 120080.

Koyano, F., Okatsu, K., Kosako, H., Tamura, Y., Go, E., Kimura, M., Kimura, Y., Tsuchiya, H., Yoshihara, H., Hirokawa, T., et al. (2014). Ubiquitin is phosphorylated by PINK1 to activate parkin. Nature 510, 162–166.

Koyle, M.L., Veloz, M., Judd, A.M., Wong, A.C., Newell, P.D., Douglas, A.E., and Chaston, J.M. (2016). Rearing the Fruit Fly *Drosophila melanogaster* Under Axenic and Gnotobiotic Conditions. J Vis Exp.

Lee, K.A., Kim, B., You, H., and Lee, W.J. (2015). Uracil-induced signaling pathways for DUOX-dependent gut immunity. Fly (Austin) 9, 115–120.

Ma, D., Storelli, G., Mitchell, M., and Leulier, F. (2015). Studying host-microbiota mutualism in *Drosophila*: Harnessing the power of gnotobiotic flies. Biomed J 38, 285–293.

Manzanillo, P.S., Ayres, J.S., Watson, R.O., Collins, A.C., Souza, G., Rae, C.S., Schneider, D.S., Nakamura, K., Shiloh, M.U., and Cox, J.S. (2013). The ubiquitin ligase parkin mediates resistance to intracellular pathogens. Nature 501, 512–516.

Marsh, J.L., and Thompson, L.M. (2006). *Drosophila* in the study of neurodegenerative disease. Neuron 52, 169–178.

Martinat, C., Shendelman, S., Jonason, A., Leete, T., Beal, M.F., Yang, L., Floss, T., and Abeliovich, A. (2004). Sensitivity to oxidative stress in DJ-1-deficient dopamine neurons: an ES-derived cell model of primary Parkinsonism. PLoS Biol 2, e327.

Mayer, E.A., Savidge, T., and Shulman, R.J. (2014). Brain-gut microbiome interactions and functional bowel disorders. Gastroenterology 146, 1500–1512.

Meissner, C., Lorenz, H., Weihofen, A., Selkoe, D.J., and Lemberg, M.K. (2011). The mitochondrial intramembrane protease PARL cleaves human Pink1 to regulate Pink1 trafficking. J Neurochem 117, 856–867.

Meulener, M., Whitworth, A.J., Armstrong-Gold, C.E., Rizzu, P., Heutink, P., Wes, P.D., Pallanck, L.J., and Bonini, N.M. (2005). *Drosophila* DJ-1 mutants are selectively sensitive to environmental toxins associated with Parkinson’s disease. Curr Biol 15, 1572–1577.

Müller-Rischart, A.K., Pilsl, A., Beaudette, P., Patra, M., Hadian, K., Funke, M., Peis, R., Deinlein, A., Schweimer, C., Kuhn, P.H., et al. (2013). The E3 ligase parkin maintains mitochondrial integrity by increasing linear ubiquitination of NEMO. Mol Cell 49, 908–921.

Myllymäki, H., Valanne, S., and Rämet, M. (2014). The *Drosophila* imd signaling pathway. J Immunol 192, 3455–3462.

Narendra, D.P., Jin, S.M., Tanaka, A., Suen, D.F., Gautier, C.A., Shen, J., Cookson, M.R., and Youle, R.J. (2010). PINK1 is selectively stabilized on impaired mitochondria to activate Parkin. PLoS Biol 8, e1000298.

Nenci, A., Becker, C., Wullaert, A., Gareus, R., van Loo, G., Danese, S., Huth, M., Nikolaev, A., Neufert, C., Madison, B., et al. (2007). Epithelial NEMO links innate immunity to chronic intestinal inflammation. Nature 446, 557–561.

Park, J., Lee, S.B., Lee, S., Kim, Y., Song, S., Kim, S., Bae, E., Kim, J., Shong, M., Kim, J.M., et al. (2006). Mitochondrial dysfunction in *Drosophila PINK1* mutants is complemented by *parkin*. Nature 441, 1157–1161.

Pesah, Y., Pham, T., Burgess, H., Middlebrooks, B., Verstreken, P., Zhou, Y., Harding, M., Bellen, H., and Mardon, G. (2004). *Drosophila parkin* mutants have decreased mass and cell size and increased sensitivity to oxygen radical stress. Development 131, 2183–2194.

Pickrell, A.M., and Youle, R.J. (2015). The roles of PINK1, parkin, and mitochondrial fidelity in Parkinson’s disease. Neuron 85, 257–273.

Poole, A.C., Thomas, R.E., Andrews, L.A., McBride, H.M., Whitworth, A.J., and Pallanck, L.J. (2008). The PINK1/Parkin pathway regulates mitochondrial morphology. Proc Natl Acad Sci U S A 105, 1638–1643.

Qin, J., Li, R., Raes, J., Arumugam, M., Burgdorf, K.S., Manichanh, C., Nielsen, T., Pons, N., Levenez, F., Yamada, T., et al. (2010). A human gut microbial gene catalogue established by metagenomic sequencing. Nature 464, 59–65.

Ren, C., Webster, P., Finkel, S.E., and Tower, J. (2007). Increased internal and external bacterial load during *Drosophila* aging without life-span trade-off. Cell Metab 6, 144–152.

Rutschmann, S., Jung, A.C., Zhou, R., Silverman, N., Hoffmann, J.A., and Ferrandon, D. (2000). Role of *Drosophila* IKK gamma in a toll-independent antibacterial immune response. Nat Immunol 1, 342–347.

Sampson, T.R., Debelius, J.W., Thron, T., Janssen, S., Shastri, G.G., Ilhan, Z.E., Challis, C., Schretter, C.E., Rocha, S., Gradinaru, V., et al. (2016). Gut Microbiota Regulate Motor Deficits and Neuroinflammation in a Model of Parkinson’s Disease. Cell 167, 1469–1480 e1412.

Sarraf, S.A., Raman, M., Guarani-Pereira, V., Sowa, M.E., Huttlin, E.L., Gygi, S.P., and Harper, J.W. (2013). Landscape of the PARKIN-dependent ubiquitylome in response to mitochondrial depolarization. Nature 496, 372–376.

Scheperjans, F., Aho, V., Pereira, P.A., Koskinen, K., Paulin, L., Pekkonen, E., Haapaniemi, E., Kaakkola, S., Eerola-Rautio, J., Pohja, M., et al. (2015). Gut microbiota are related to Parkinson’s disease and clinical phenotype. Mov Disord 30, 350–358.

Sharon, G., Sampson, T.R., Geschwind, D.H., and Mazmanian, S.K. (2016). The Central Nervous System and the Gut Microbiome. Cell 167, 915–932.

Shiba-Fukushima, K., Imai, Y., Yoshida, S., Ishihama, Y., Kanao, T., Sato, S., and Hattori, N. (2012). PINK1-mediated phosphorylation of the Parkin ubiquitin-like domain primes mitochondrial translocation of Parkin and regulates mitophagy. Sci Rep 2, 1002.

Shiba-Fukushima, K., Inoshita, T., Hattori, N., and Imai, Y. (2014). PINK1-mediated phosphorylation of Parkin boosts Parkin activity in *Drosophila*. PLoS Genet 10, e1004391.

Shin, S.C., Kim, S.H., You, H., Kim, B., Kim, A.C., Lee, K.A., Yoon, J.H., Ryu, J.H., and Lee, W.J. (2011). *Drosophila* microbiome modulates host developmental and metabolic homeostasis via insulin signaling. Science 334, 670–674.

Taira, T., Saito, Y., Niki, T., Iguchi-Ariga, S.M., Takahashi, K., and Ariga, H. (2004). DJ-1 has a role in antioxidative stress to prevent cell death. EMBO Rep 5, 213–218.

Téfit, M.A., and Leulier, F. (2017). *Lactobacillus plantarum* favors the early emergence of fit and fertile adult *Drosophila* upon chronic undernutrition. J Exp Biol 220, 900–907.

Wang, Q., Garrity, G.M., Tiedje, J.M., and Cole, J.R. (2007). Naive Bayesian classifier for rapid assignment of rRNA sequences into the new bacterial taxonomy. Appl Environ Microbiol 73, 5261–5267.

Wong, A.C., Chaston, J.M., and Douglas, A.E. (2013). The inconstant gut microbiota of *Drosophila* species revealed by 16S rRNA gene analysis. ISME J 7, 1922–1932.

Wong, A.C., Luo, Y., Jing, X., Franzenburg, S., Bost, A., and Douglas, A.E. (2015). The Host as the Driver of the Microbiota in the Gut and External Environment of *Drosophila melanogaster*. Appl Environ Microbiol 81, 6232–6240.

Wong, M.L., Inserra, A., Lewis, M.D., Mastronardi, C.A., Leong, L., Choo, J., Kentish, S., Xie, P., Morrison, M., Wesselingh, S.L., et al. (2016). Inflammasome signaling affects anxiety- and depressive-like behavior and gut microbiome composition. Mol Psychiatry 21, 797–805.

